# Improving and evaluating deep learning models of cellular organization

**DOI:** 10.1101/2022.05.24.493229

**Authors:** Huangqingbo Sun, Xuecong Fu, Serena Abraham, Jin Shen, Robert F Murphy

## Abstract

**Motivation:** Cells contain dozens of major organelles and thousands of other structures, many of which vary extensively in their number, size, shape and spatial distribution. This complexity and variation dramatically complicates the use of both traditional and deep learning methods to build accurate models of cell organization. Most cellular organelles are distinct objects with defined boundaries that do not overlap, while the pixel resolution of most imaging methods is not sufficient to resolve these boundaries. Thus while cell organization is conceptually object-based, most current methods are pixel-based. Using extensive image collections in which particular organelles were fluorescently-labeled, deep learning methods can be used to build conditional autoencoder models for particular organelles. A major advance occurred with the use of a U-net approach to make multiple models all conditional upon a common reference, unlabeled image, allowing the relationships between different organelles to be at least partially inferred.

**Results:** We have developed improved GAN-based approaches for learning these models and have also developed novel criteria for evaluating how well synthetic cell images reflect the properties of real images. The first set of criteria measure how well models preserve the expected property that organelles do not overlap. We also developed a modified loss function that allows retraining of the models to minimize that overlap. The second set of criteria uses object-based modeling to compare object shape and spatial distribution between synthetic and real images. Our work provides the first demonstration that, at least for some organelles, deep learning models can capture object-level properties of cell images.

**Availability:** A Reproducible Research Archive containing all source code, generated images and analysis results will be made available at http://murphylab.cbd.cmu.edu/Software upon publication.

**Supplementary information:** Supplementary data are available at *Bioinformatics* online.

## 1 Introduction

Deep learning been very successfully used in recent years for biomedical image analysis applications, including to analysis and modeling of fluorescence microscope images. Fluorescence microscopy is a fundamental and powerful approach in the life sciences that is used to observe cells and their structures by labeling them with fluorophores or fluorophore-conjugated antibodies [Lichtman and Conchello, 2005]. Deep learning applications to fluorescence microscope images have included reconstruction of super-resolution images [Ouyang *et al*., 2018], cell segmentation [Stringer *et al*., 2021, Greenwald *et al*., 2021], integrative tissue analysis [Maric *et al*., 2021, Bao *et al*., 2022] and various augmented microscopy approaches [Wang *et al*., 2021].

Despite its power, there are significant limitations of fluorescence microscopy for observing and modeling the complex variation in number, size, shape and spatial distribution of many organelles and other subcellular structures. One is the limited spatial resolution that makes it difficult to resolve individual organelles. Another is the limited number of fluorescent tags that can be observed simultaneously. While this can be partially overcome by multiplexing in fixed cells [Schubert *et al*., 2006, Lin *et al*., 2015, Goltsev *et al*., 2018], it limits learning of complex relationships between multiple structures in live cells.

Nonetheless, learning and modeling these relationships is an important challenge in cell biology. Initial work using traditional computer vision methods introduced the idea of building generative models of cells in which organelle positions within cells EW conditional upon other parts, such as the cell membrane, nuclear membrane and microtubules [Zhao and Murphy, 2007, Johnson *et al*., 2015, Majarian *et al*., 2019]. With the advent of deep learning and the creation of large collections of images labeled for specific organelles, a significant advance occurred through the creation of autoencoder models for organelles that were also conditional on cell and nuclear membranes [Johnson *et al*., 2017, Donovan-Maiye *et al*., 2022].

This conditional approach has been taken even further by constructing deep learning models that are conditional upon an easily acquired common reference image, such as a transmitted light image. In this “in silico labeling” or “label-free microscopy” approach, separate deep neural networks are trained to predict the likely distribution of specific organelle markers from transmitted light or other label-free images. Christiansen *et al*. [2018] first proposed this method to label the nucleus and neurons; Ounkomol *et al*. [2018], Cooke *et al*. [2021], Waibel *et al*. [2019] dramatically extended this approach by in silico labeling of multiple subcellular structures. These approaches used variations on the powerful U-Net model with a series of pooling and up-sampling operations that helps the model to capture image information at different scales [Ronneberger *et al*., 2015, Falk *et al*., 2019]. Wang *et al*. [2021] described an improved U-Net architecture which inserted a self-attention module into the U-Net down-sample module to help enlarge the receptive field of the model. Nonetheless, as with other autoencoder-based models, these implementations are trained with only a pixel wise loss function (like mean squared error, MSE) which often results in their producing relatively blurred images [Huang *et al*., 2018]. These may not accurately reflect the edge morphology of subcellular structures, especially for smaller objects. One alternative is to modify the convolutional neural network (CNN) architectures. Deep Recurrent Attentive Writer (DRAW) [Gregor *et al*., 2015] was proposed to generate realistic images using a variational autoencoder (VAE) with recurrent blocks. The subsequent convolutional DRAW [Gregor *et al*., 2016] further combined the recurrent blocks with convolution components to improve the model. Another alternative is to modify the loss function used in training. PixelRNN and PixelCNN are well-known generative models, which employ an auto-regressive method to learn the explicit distribution [Van Oord *et al*., 2016, Van den Oord *et al*., 2016, Salimans *et al*., 2017]. As a last alternative, Generative Adversarial Network (GAN) has been shown to generate natural, realistic instances [Goodfellow *et al*., 2014]. Unlike traditional CNN models trained with a pixel-wise loss function, GANs have two components: a generator and a discriminator. The generator is trained to produce instances that is similar to the instances from the target domain, and the discriminator will distinguish whether an instance is a real or generated instance. Using an adversarial training loss, the blurry instances can be easily recognized as generated instances, forcing the generator to produce sharper and more natural instances.

“Pix2Pix” network is a GAN variant that combines a U-Net-based generator and a PatchGAN discriminator [Isola *et al*., 2017]. Experiments show that the Pix2Pix network performs well on image-to-image translation. Predicting fluorescence or stain distributions is also an image-to-image translation problem, and prior work such as Rivenson *et al*. [2019] has used a variant of the pix2pix network for virtual tissue staining.

While previous work has shown promising results, there is significant room for improvement, especially for in silico labeling of multiple cellular structures. One potential direction is to design more efficient DL model architectures, and another is to set up novel model-training approaches and objectives specialized for this task. The current state-of-the-art approaches treat each protein channel (subcellular structure) independently by building separate models from the reference input. These essentially give the probability of each pixel being in a given structure (or equivalently the amount of each structure expected to be present in each pixel), but do not address any limitations on whether more than one structure can be in the same pixel. In other words, they do not address the concept of exclusivity of structures, which, while complicated by the pixel resolution, is expected for most membrane-bound organelles. Depending on pixel resolutions, a single pixel may cover boundaries of two organelles. Nonetheless, a low fraction of pixels with overlapped organelles is still expected. Further, most current supervised learning-based computer vision approaches applied in biological studies only rely on pixel-wise evaluations, which does not consider the number and shape of individual biological components nor their spatial distributions. Also, while there is a large body of work on designing generative models, there is only limited work focused on criteria for evaluating generative models in general [Szegedy *et al*., 2016, Barratt and Sharma, 2018]. Therefore, higher-order metrics can be useful to better understand the quality of generative models.

In this work, we first describe an improved 3D Pix2Pix network with recurrent residual convolutional units [Alom *et al*., 2018]. Without increasing the size of the network, recurrent units allow a deeper network. The residual-like block also gains the advantage of ResNet to maintain the deep network [He *et al*., 2016]. Then, we proposed novel evaluation metrics for multi-channel reconstructed fluorescence images. The first we define as “exclusivity”. This criterion describes the self-consistency of a predicted image containing multi-subcellular organelles under the assumption that overlap between those organelles should be minimal. As a result, no strong fluorescence signal is expected from multiple channels at one single pixel. Note that a single image shows only one fluorescence-tagged organelle, which means that we cannot directly compare different organelle patterns. Therefore we trained the original models for all organelle and made predictions of all protein channels for every single transmitted light image for simultaneous evaluation of the exclusivity among different organelles. Also, we showed efficient ways to improve the exclusivity of in silico labeled images by using both modified loss functions with an additional “exclusivity” term and novel neural network architectures compared to the previous work by Ounkomol *et al*. [2018]. We also propose novel metrics for object-wise evaluations of subcellular structures. In particular, we compare object shapes between real and synthetic images by using spherical harmonic parameterization, which has been empirically proven to be efficient and robust on object’s representation [Ruan and Murphy, 2019, Styner *et al*., 2006, Viana *et al*., 2021]. An overview of the approaches used for evaluating and improving generative models of organelles is provided in Figure 1.

**Fig. 1.**
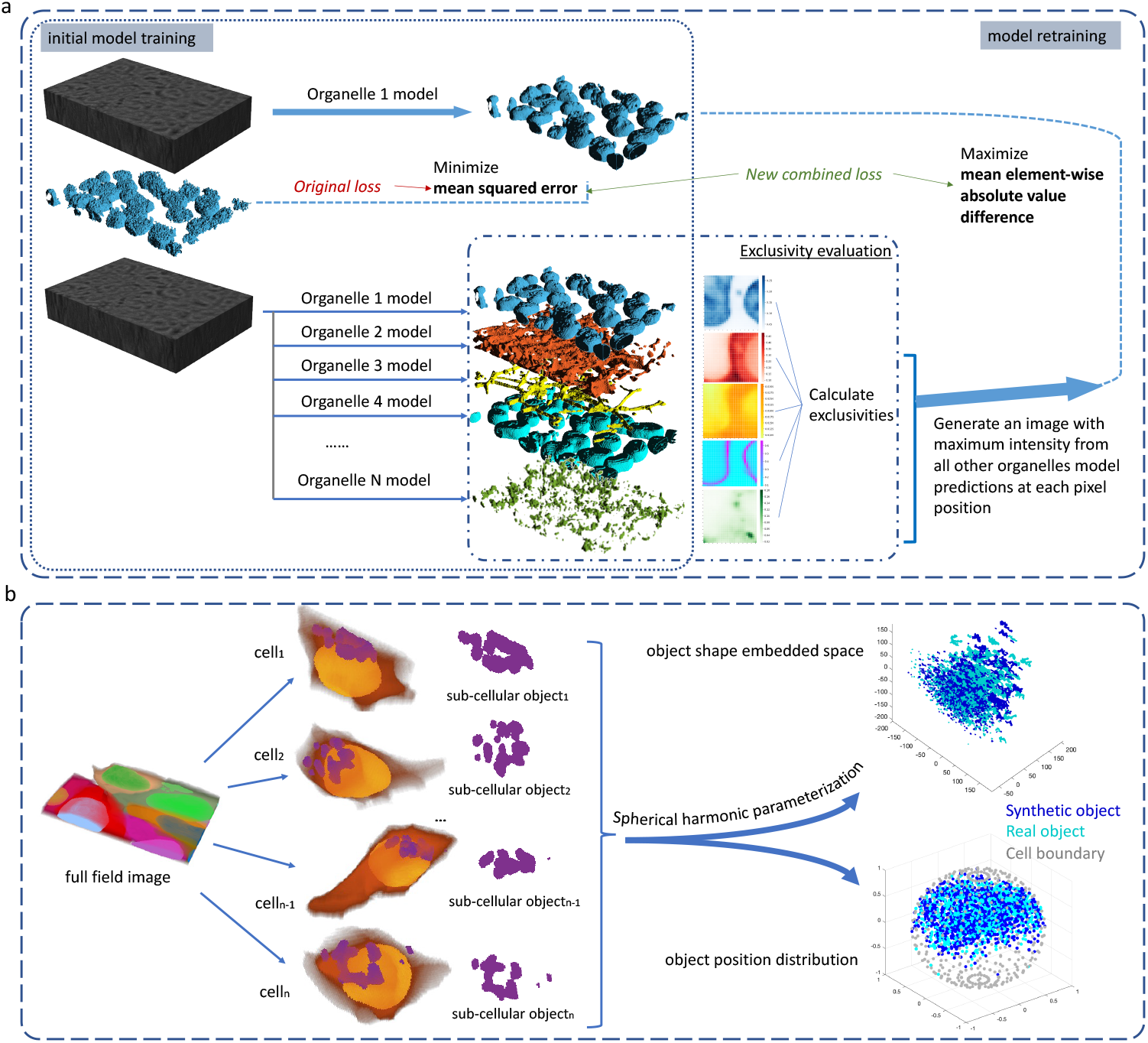
Approaches to evaluating and improving organelle generative models. (a) The model training process described by Ounkomol et al. [2018] is shown on the left and the process of evaluating overlap between predicted organelles and retraining models to reduce it is shown on the left. After training individual organelles models, each model is retrained in sequence using the additional loss term that seeks to minimize overlap (maximize exclusivity). (b) The process for object-based evaluation of synthetic images using CellOrganizer is illustrated. For each organelle model, real and synthetic image are segmented into individual objects. The real and synthetic object shapes are described using spherical harmonic parameterization and displayed in a 3-dimensional embedding, and their positions normalized to a spherical cell are also shown. The distributions of real and synthetic objects in both shape and position are compared statistically as described in the Methods.

## 2 Methods

### 2.1 Image dataset

We performed our evaluations using the image dataset, created by the Allen Institute for Cell Science, that was used by Ounkomol *et al*. [2018] (available from https://downloads.allencell.org/publication-data/label-free-prediction/index.html). It contains fluorescence microscope images for 12 different subcellular components (actin filaments, microtubules, endoplasmic reticulum, desmosomes, cell membrane, actomyosin bundles, Golgi apparatus, DNA, nuclear envelope, mitochondria, nucleoli, and tight junctions) and their corresponding transmitted light images. For each subcellular component, we divided the dataset into 60 images for training and 20 images for testing.

### 2.2 Deep neural network architecture and training and validation

We implemented three deep neural network architectures for training organelle models. The first is the same as the one proposed in Ounkomol *et al*. [2018], which we will refer to as the U-Net model for simplicity. For each organelle, the U-Net model was trained for 100000 iterations.

The second is a model adapted from the original Pix2pix network model, which we refer to as a Vox2Vox network with recurrent units (Vox2Vox-RU). This model contains two parts: a discriminator with a stack of 3D convolutional layers; a generator modified from the 3D version of recurrent residual convolutional neural network based on U-Net (R2U-Net) [Alom *et al*., 2018]. The essential idea in this model is to replace the convolutional layers by recurrent residual convolutional units (RRCU) in the U-Net. RRCU consists of a 1 × 1 convolution layer and 2 subsequent recurrent convolutional layers (see Figure 3 and 4 in Ounkomol *et al*. [2018]), which increases the model depth without adding new parameters but also adopts the idea from ResNet to mitigate limitations of deep net training. The detailed architecture of our model can be found in Figure 2.

**Fig. 2.**
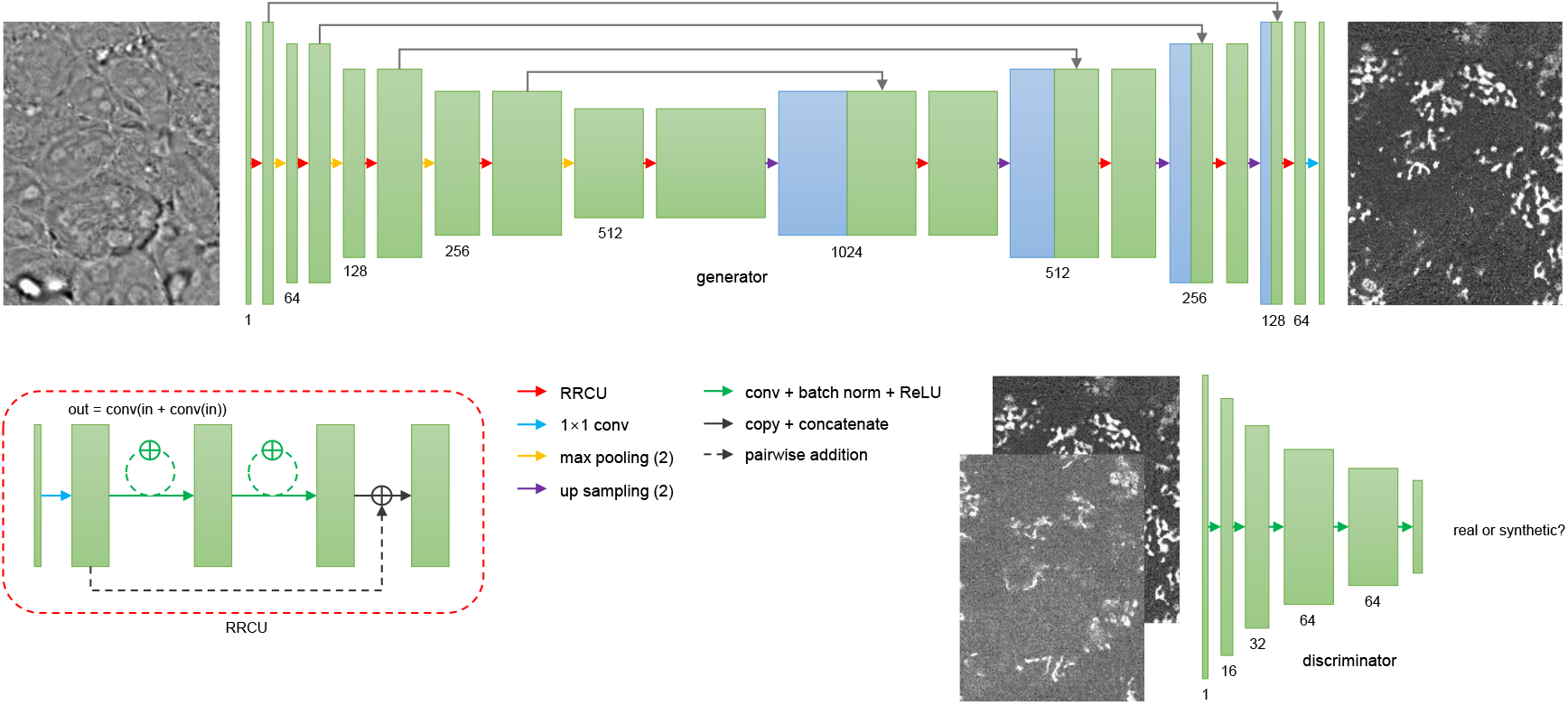
Neural network architecture of our proposed Vox2Vox-RU network.

**Fig. 3.**
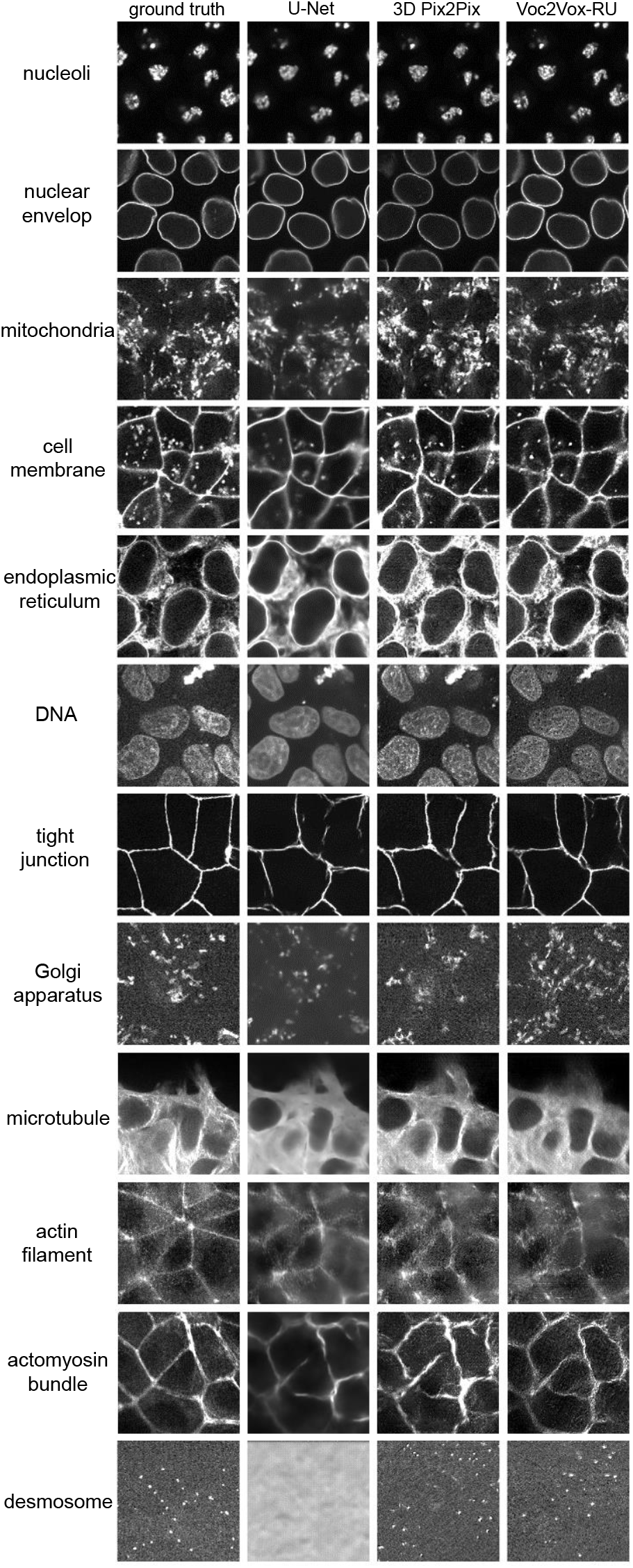
Example real and predicted images.

**Fig. 4.**
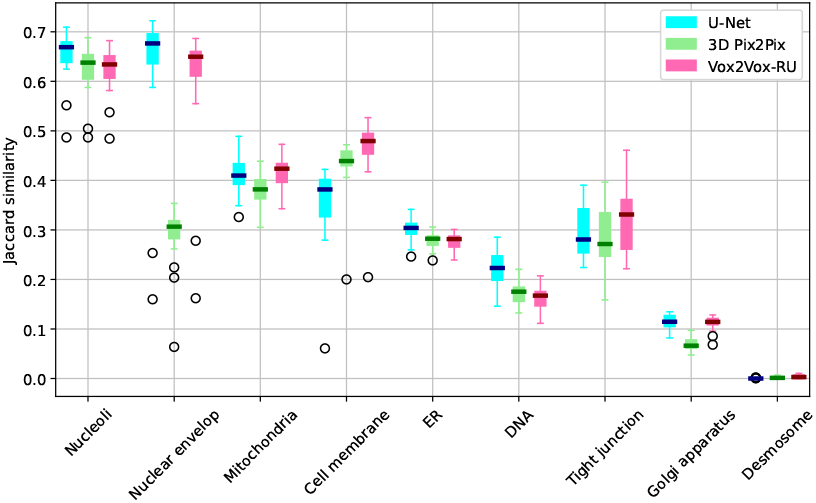
Image-wise Jaccard similarity are calculated based on segmented images for both models comparing to the segmented real images using the segmentation tool provided by Chen et al. [2020]. The blue bars refer to the U-Net model. The green bars refer to the 3D-Pix2Pix model and the pink bars refer to the Vox2Vox-RU model. The short line in the center of each box refers to the median value.

The third is a 3D Pix2Pix model whose generator is the same as the U-Net and whose discriminator is the same as shown in Figure 2.

Both 3D Pix2Pix and Vox2Vox-RU models were trained for a total of 20000 iterations with an Adam optimizer with a learning rate of 2 × 10^*−*4^ and a batch size of 4. To provide a “warm start” in order to maintain stable training, for the first 2500 iterations the value of real and fake labels provided for the discriminators were set to

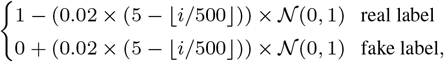

where *i* refers to the iteration number. The soft labels prevent the discriminator from learning much faster than the generator which can lead the network to fail to converge.

After initial training of all organelle models by both methods, we predicted all organelle signals with the trained 12 models for all transmitted light images for both training approaches.

### 2.3 Evaluation of organelle exclusivity

As discussed in the Results, we propose that good organelle models should have a large difference between the predicted amount of the “major” organelle and that of other organelles for each pixel in an image. We therefore defined three descriptions for this “exclusivity”. Overall exclusivity was defined as the difference between the highest predicted value and the second highest predicted value, averaged over all pixels. Organelle exclusivity was defined for each organelle individually as the average for all pixels for which that organelle has the highest predicted value of the difference between that predicted value and the second highest predicted value. Pairwise exclusivity was defined for each directed pair of organelles as the average for all pixels for which the first organelle has the highest predicted value of the difference between that predicted value and the predicted value for the other organelle (thus the pairwise exclusivity is not symmetric).

### 2.4 Retraining

After training the original prediction, we proposed a new loss function, making adjustments to the current model to enlarge the exclusivity. We retrained all the models with the reverse order of the evaluated organelle exclusivity. The new loss function consists of two part, one is to minimize the original mean squared error (MSE) for U-Net model and GAN loss for Vox2Vox-RU model, the other is a new term which maximizes the mean element-wise absolute value difference between the prediction of the current model and the maximum intensity at each pixel position of all other subcellular component predictions. The second term aims to increase the exclusivity of each pixel of the trained models. For example, to retrain a model for ER prediction, given 12 trained models for different subcellular components, if ER has the highest intensity at the pixel position, then we want to enlarge the differences between the pixel intensity of ER and the second highest pixel intensity of among the rest of 11 subcellular components to increase the exclusivity; if ER is not the highest intensity at the pixel position, then we want it to be as small as possible compared with the subcellular component with the largest intensity to increase the exclusivity.

The U-Net model was fine-tuned with 12500 additional iterations for retraining. We used a parameter, *p*, to control the extent of maximizing the exclusivity in the loss function. We scanned through a potential list of candidate values of *p* and used the criteria of 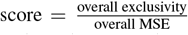 to select the parameter with the highest score for each model. We update the predictions after each subcellular component model was retrained, based on which the next subcellular component was retrained.

For retraining the Vox2Vox-RU model, to better balance three loss terms (GAN loss, MSE/L1 loss, exclusivity loss) and save the computational effort, the weight of exclusivity loss in Vox2Vox-RU is mannually set.

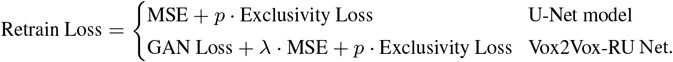

The loss function for training Vox2Vox-RU has an additional L2/L1 loss term (differs from different subcellular components) between the prediction and real fluorescence tag which helps to further stablize the training process, the detailed loss functions used are reported in Table S1. The different loss function terms are mannually set mainly based on the texture of subcellular patterns. For example, for tubule-like structure, since the training image resolution is limited (which makes tubule-like structure patterns blurry), we prefer using L2-loss; for membrane-like structures, to maintain a thin reconstructed structure, we prefer using L1-loss.

Unlike U-Net model retraining, Vox2Vox-RU was retrained from scratch with the same optimizer for 25000 iterations, still with a warm start.

### 2.5 Subcellular object shape and spatial distribution analysis

Spherical harmonic transform uses a series of orthogonal functional to fit the surface of a genus-0 topology object, which provides an efficient way to parameterize the objects with spherical harmonic descriptors [Nain *et al*., 2007, Ruan and Murphy, 2019]. The quality of spherical harmonic parameterization can be assessed by calculating the Hausdorff distance between the original shape’s surface and the reconstructed shape’s surface. The quality of spherical harmonic parameterization heavily depends on the order of spherical harmonics; we set it to 31 in this work.

To compare the synthetic images from DL models and real images, the objects were first segmented using the segmentation tool provided by Chen *et al*. [2020]. In order to examine the subcellular components’ spatial distribution in the cell, we also performed the segmentation for the cell nucleus and cell boundary in the images. Then, the partial cells at the boundary of the images as well as the objects in those cells were discarded. The valid objects were parameterized via spherical harmonic transform. To compare if the objects from the synthetic images are comparable in spherical harmonic descriptor space, principle component analysis is performed to reduce the dimensionality of the spherical harmonic descriptor to be smaller, we make it as 3 in this work. With two population pf low-dimensional shape descriptors *x*_1_, …, *x*_*m*_, *x*_*m*+1_, …, *x*_*m*+*n*_, where *m* denotes as the number of objects in synthetic images and *n* denotes as the number of objects in real images, we also give them labels to indicate the population coming from as *y*_1_ …, *y*_*m*_ = 1 and *y*_*m*+1_, …, *y*_*n*_ = 2. Inspired by the k-clique percolation approach [Palla *et al*., 2005], for each object, we find its *k* (*k* = 8 in this work) nearest neighbors and check the purity of labels in its neighbor objects. Ideally, if two population merges well, the label fraction will be close to 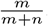, and if two population lay separately in the spherical harmonic descriptor space, the purity of neighbor objects will be higher. Formally, we calculate

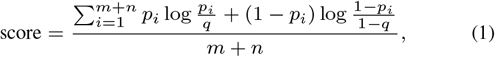

which is the average KL-divergence between the neighbor objects purity 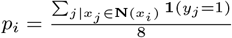 and reference purity 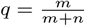, where **N**(·) returns the 8-nearest neighbor object set and **1**(·) refers to the indicate function.

To compare the spatial distribution of subcellular components inside the cell, we first align all valid cells by their major axis, then map the cell boundary to a unit sphere and the nucleus to the center of the sphere. All the centers of objects in the cell are mapped onto these polar coordinates, and a Gaussian kernel is used to generate a continuous object distribution in the unit sphere domain. The KL-divergence was then calculated between this fit for synthetic images and real images.

This analysis of organelle shape and spatial distribution was performed using version 2.9.3 of the open source CellOrganizer system (http://CellOrganizer.org, https://github.com/murphygroup/cellorganizer).

## 3 Results

### 3.1 Comparing organelle model learning approaches

We began by training individual organelle models with U-net, 3D Pix2Pix and our new approach. We used the same dataset described by Ounkomol *et al*. [2018], and randomly divided it into training and test sets. Figure 3 shows a comparison between predictions from the U-Net, 3D Pix2Pix and Vox2Vox-RU models for nine different organelles. When examined visually, all three of the models give reasonable predictions compared to the ground-truth, although the 3D Pix2Pix and Vox2Vox-RU images are closer to the ground truth in terms of preservation of contrast level, detailed texture, and sharpness of organelle boundaries. This difference is most pronounced for the desmosome model. When evaluated by the overall mean-square error (MSE) between the original and predicted images, most U-Net models perform better than 3D Pix2Pix and Vox2Vox-RU models (Table S2). However, the U-Net desmosome model has a very high MSE, as may be expected given the visual difference. The 3D Pix2Pix and Vox2Vox-RU models perform reasonably for all organelles such that their overall MSE is lower, but this is reversed when the desmosome model is removed from the averages. These results are consistent with the difference in goals of the two approaches U-Net and GAN-based models: the GAN-based models will produce models with better “high level” performance at the sacrifice of a small increase in MSE.

We therefore sought to quantitatively evaluate the performance of the three modeling approaches from the “high level” viewpoint of subcellular organelle morphology. To do this, we performed segmentations of original and synthetic images following the approach described by Chen *et al*. [2020]. We then calculated the Jaccard similarity between the segmentations for all three approaches (Figure 4). We observed that the segmentation results for tubular organelles (microtubule, actin-filaments and actomyosin) were poor, and therefore did not include these organelles in the comparison.

All three modeling approaches performed similarly for most of the organelles. Both U-Net and Vox2Vox-RU performed best for the relatively easy task of generating nuclear components (nucleoli and nuclear envelope), whereas 3D Pix2Pix had comparable results for nucleoli but poor results for nuclear envelope. It is worth noting that Vox2Vox-RU significantly outperforms U-Net in cell membrane prediction, with a roughly 10% higher median value and 3D-Pix2Pix falls in between; this is consistent with the observation in Figure 3 that the U-Net model produces thicker cell membranes than those in the original image. Also for nuclear envelope, mitochondria and Golgi apparatus, 3D Pix2Pix has inferior performance compared with other two models, and has never achieved top performance for any organelle, which may indicate that changes in Vox2Vox-RU is useful for GAN model and may even negatively affect the performance without them for GAN model. Note that these results evaluate *semantic* segmentation which simply seeks to distinguish organelle foreground from cell background and does not consider *instance* segmentation which evaluates individual organelle morphology. We will consider this later. Also, the Vox2Vox-RU outperforms 3D Pix2Pix in Jaccard similarity on 6 out of 9 organelles and almost ties on the rest of 3 organelles. This suggests that GAN-based training as well as the changes in the generator of Vox2Vox-RU contribute to the improvement.

### 3.2 Exclusivity analysis and retraining

Both the training and retraining processes of U-Net and Vox2Vox-RU models for each organelle are carried out separately, and therefore, when the resulting models are used to make predictions for a given transmitted light image, their predictions are independent. In other words, an intensity level is predicted for each pixel for each organelle, and it is entirely possible that high intensity values will be predicted for the same pixel for more than one organelle. Such “overlap” between organelles in the same pixel can result from the two organelles having similar characteristics in transmitted light images, or from the learned models having significant blur or uncertainty around the edges of predicted organelles. The first may be inherent in the approach, but the second can be a reflection of the quality of the training. We therefore propose new criteria for evaluation of label-free microscopy - organelle “exclusivity”. The premise is that most pixels in an image should have only one organelle with high predicted content. This concept can be quantified for all organelles, each organelle individually relative to all others, and each organelle relative to each other organelle (Figure 5).

**Fig. 5.**
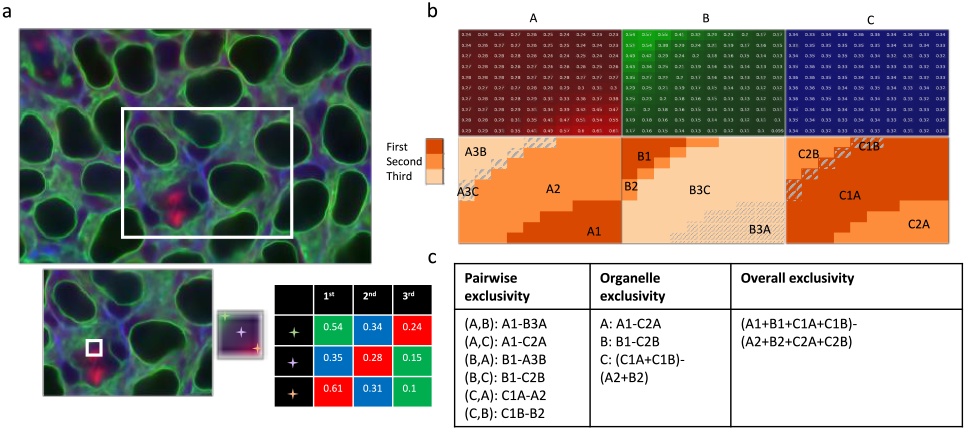
An illustration of different metrics for exclusivity. (a) An example of a combined image showing the values of three organelle predictions using red, green and blue. The insets show magnification of smaller and smaller regions. (b) Predictions for the smallest region in (a). The first row shows the intensity of each organelle channel (A,B,C). The second row shows the pixel-wise intensity relationships. The darkest orange indicates that the channel in that column has the highest intensity among all three at that position, the lighter orange color indicates that the channel has the second highest intensity, and the lightest color indicates that the channel has the lowest intensity. The code in each subregion summarizes its properties: the first letter and number indicate the ranking of that channel in that subregion, and the second letter (if present) indicates which channel is highest in that subregion. (c) A detailed example of the definitions for the different exclusivity measure. The use of a code in the definitions represents the pixel-wise operations corresponding to that code (“-” is pairwise subtraction and “+” is pairwise addition).

The organelle-specific exclusivities for all three models, as well as the overall exclusivities, are reported in Table 1. Our proposed Vox2Vox-RU model reaches a higher overall exclusivity compared with U-Net, and 3D Pix2Pix falls in between. Some organelles such as nucleoli and nuclear envelopes have high exclusivity and low MSE for both U-Net and Vox2Vox-RU (Table S2). On the other hand, microtubules, actin filaments, and endoplasmic reticulum have both low MSE and low organelle exclusivity, suggesting that despite the accuracy of the prediction compared with the true protein signals, the models are limited in their ability to distinguish between them. As noted above, the desmosomes U-Net model performs poorly in MSE due presumably to their small sizes, yet the Vox2Vox-RU and 3D Pix2Pix model is still able to produce predictions with relatively reasonable MSE. All desmosome models have higher exclusivity than microtubules, actin filaments, and ER, indicating that they are better distinguished than those organelles.

**Table 1.** The organelle exclusivity (See Figure 5 for the defination) on the test set of the initial U-Net 3D Pix2Pix, and Vox2Vox-RU models. Note that exclusivity values are in units of pixel intensities.

|  | U-Net | 3D Pix2Pix | Vox2Vox-RU |
| --- | --- | --- | --- |
| microtubule | 0.409 | 0.560 | 0.57 |
| actin filament | 0.401 | 0.527 | 0.577 |
| desmosome | 0.526 | 0.61 | 0.659 |
| DNA | 1.05 | 0.892 | 0.86 |
| nucleoli | 1.50 | 1.931 | 1.69 |
| nuclear envelope | 1.11 | 0.966 | 1.11 |
| cell membrane | 0.756 | 1.009 | 1.05 |
| actomyosin bundle | 0.738 | 1.003 | 0.807 |
| endoplasmic reticulum | 0.433 | 0.565 | 0.58 |
| Golgi apparatus | 0.774 | 0.698 | 0.767 |
| mitochondria | 1.41 | 1.180 | 1.2 |
| tight junction | 1.50 | 1.007 | 1.04 |
| Overall | 0.7930 | 0.847 | 0.8635 |

The overall exclusivity of trained 3D Pix2Pix model is at the middle place (0.8470), which further shows that both GAN training and the architecture change in generator are needed. Comparing the organelle exclusivity values with the mean square errors, we see little correlation for the U-net models (R2=-0.03) and mild negative correlation for the Vox2Vox-RU models (R2=-0.38). This reinforces the conclusion from visual analysis that the GAN-training can capture both aspects, and strongly indicates the value of not evaluating organelle model methods using MSE alone.

In view of the large overlap (low exclusivity) for some organelle models, we next explored whether we could retrain the models to increase the exclusivity. To do this, we added an additional term to the loss function of either modeling approach so that learning would balance reducing MSE and increasing exclusivity for both U-Net and Vox2Vox-RU. We used a greedy approach to improve overall exclusivity, retraining each organelle model in the order of decreasing exclusivity. However, a critical issue was how to weight the contribution of exclusivity to the overall loss. As described in the Methods, for the U-Net model, we performed a grid search to determine the optimal weight of the exclusivity loss term, *p* (Figure 6). This was chosen for each organelle in turn by optimizing the ratio of the resulting overall exclusivity and MSE; the model for that organelle was then replaced with the optimal model.

**Fig. 6.**
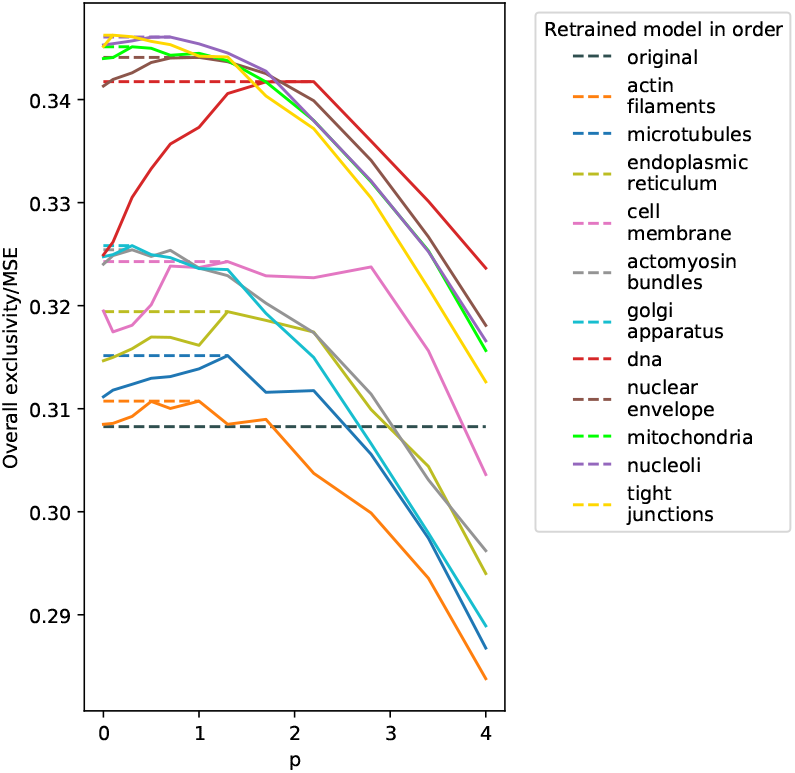
The overall improvement trajectory of the ratio of overall exclusivity and MSE through retraining of different protein models using U-Net. The weight *p* was scanned from 0 to 4 and the optimal *p* and corresponding retrained model was chosen if the overall exclusivity MSE ratio reached the maximum, based on which the next organelle model was trained.

All 12 models were retrained except desmosome (due to the extremely large MSE after the original training process compared to other organelles the retraining would be inappropriately biased towards extremely large exclusivity when using the ratio criterion). As can be seen, the overall exclusivity gradually rose as each model was retrained, and the optimal weights for the different organelles were roughly similar. When retraining the Vox2Vox-RU models, we did not do a grid search over weights but rather manually assigned a fixed weight for a given organelle; this was because of the high computational cost for grid search and potential difficulty maintaining a stable training process (see Table S1). After the whole retraining process, the overall exclusivity increased about 24% and 13.17% for U-Net and Vox2Vox-RU models on the test set. The smaller improvement on Vox2Vox-RU models could be due to the optimization of the loss term for the U-Net but not the Vox2Vox-RU models. Example fluorescence tag predictions from the retrained models are shown in Figure S1.

Comparisons of organelle exclusivity and MSE before and after the retraining for both models are shown in Figure 7. Overall, for both models, most organelles sacrifice a slight increase in MSE to get a fair increase in organelle exclusivity. The improvement in pairwise exclusivity upon retraining is generally consistent with the changes in organelle exclusivity, as shown in (Figure S2). We also record changes of each organelles exclusivity during retraining and typically see a slight decrease in exclusivity of other organelles after a specific organelle model is retrained (Figure S3).

**Fig. 7.**
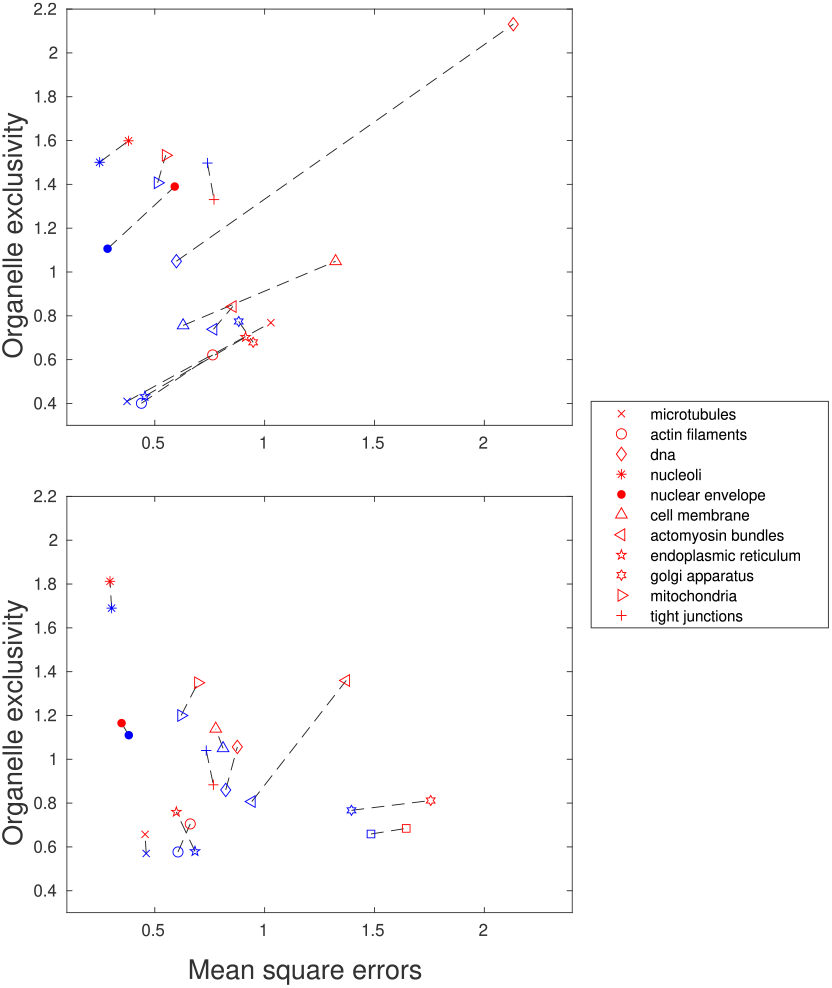
The MSE and organelle exclusivity before (blue) and after (red) the retraining for U-Net (upper) and Vox2vox (bottom), connected by dashed lines for specificity.

### 3.3 Subcellular component shape and spatial distribution analysis

The criteria we have described so far measure reconstruction error both at the level of individual pixels and at the level of segmentation into regions above and below a threshold. The exclusivity metric we have introduced sought, in part, to evaluate a different aspect, the sharpness of the boundaries between predicted organelles. Since most of the models we have evaluated are for organelles that consist of distinct objects, we next sought to evaluate whether those models could properly capture the number, shape and cellular positions of those organelles. We began by segmenting all real and synthetic images into individual cells (ignoring incomplete cells that extend beyond the boundary of the image) and then segmenting individual organelles. This yields a set of objects for each complete cell for each image for each organelle. Each object was represented by a list of the pixels contained within it (or equivalently by an individual binary image) and by the coordinates of its centroid. To evaluate the consistency between the object shapes from real and synthetic images, we chose, based upon our prior work on evaluation of models of cell and nuclear shape [Johnson *et al*., 2017], to use spherical harmonic (SPHARM) parameterization to construct object shape models. Our previous results showed that our modified implementation of SPHARM modeling performed better than other available methods (including deep learning methods) for modeling even eccentric 3D cell shapes. However, this parameterization approach only applies to genus-0 objects (those without holes), and we therefore only created object-based models for mitochondria, nucleoli, Golgi apparatus, and desmosome.

Each object is described by a high-dimensional shape vector that can be converted (back-transformed) into a shape. We can then create a generative model of object shape by choosing a lower dimensional embedding of the vectors. To confirm that each object’s shape is being accurately represented by the SPHARM parameterization, we first calculate the Hausdorff’s distance between the original object and its reconstruction from the lower dimensional embedding (Hausdorrf’s distance is the largest difference between the locations of equivalently spaced points). As shown in Table S3, most of the objects are well-fitted with average Hausdorff distance around 1 *μm*. We also observe that the number of objects produced by both models is comparable to the real images.

Given two sets of objects (e.g., from synthetic and real images), we can construct a lower-dimensional embedding of the object shapes using just the first three principal components of the SPHARM vectors of both sets. The two sets can be visually compared in plots in which downsampled versions of each shape are displayed at their position in the 3D embedding (Figures S5, S4, S6, and S7).

To make the comparison quantitative, we compared the distributions of the two sets in the reduced spherical harmonic descriptor space. As described in the Methods, the principle is that the similarity of the object distributions can be compared by measuring the fraction of the neighbors of each object that are real objects. We calculate this as the KL-divergence (eq. 1) between the observed fraction and the expected fraction based upon the number of objects from each type of image.

Unlike object shapes that can be compared directly, comparing the spatial distributions of objects within the cell requires conversion of each object’s Cartesian coordinates within the image into some common frame of reference. We chose a spherical cell as this reference and mapped each object’s centroid into polar coordinates in which the radius is the fractional distance of the object from the cell membrane. The distributions can be visually compared in Figures S9, S8, S10, and S11). To make the comparison quantitative, the probability density distributions of the objects of each type are separately fit using Gaussian kernels which allows us to compare the two probability densities directly by the KL-divergence (as described in the Methods).

Figure 8 shows a scatter plot of the two divergences versus each other, in which low values for both (the lower left corner) correspond to similar spatial and shape distributions. Both models perform well on nucleoli (especially after retraining), which can be expected since the shape of nucleoli is relatively regular and they are located in the relatively small nuclear which leads to a very small spatial distribution divergence for both models. The Vov2Vox-RU model outperforms the U-Net model on the Golgi apparatus and does slightly better on mitochondria. This is consistent with our observation in Figure 3 that Vov2Vox RU model produces objects with sharper edges and appearances more similar to the objects in the real images. In contrast, modeling the desmosome is the hardest task for both models, since it is small in size and distributed throughout the entire cellular environment. The U-Net models fails to produce reasonable desmosome objects, and the Vox2Vox-RU model produces desmosome objects which are comparable to the real objects but fails to accurately recover its spatial distribution (see the p-values in Table S4).

**Fig. 8.**
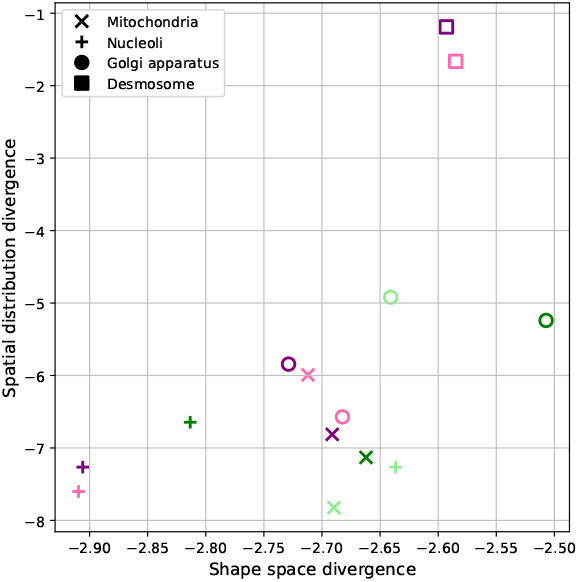
Models’ divergence in spherical harmonic descriptor shape space (x-axis) and cellular spatial distribution (y-axis). The values are reported in logarithm, and the lower left corner is ideal which indicates low divergence in both shape and spatial distribution comparing to the real objects. The pink and purple markers refer to the original and retrained Vox2Vox-RU model, respectively. The light green and dark green markers refer to the original and retrained U-Net model, respectively.

To estimate the statistical significance of the divergences, we performed a permutation test to establish the divergence expected between objects from two different sets of real images. The results and p-values are shown in Table S4 and S5. Large p-values indicate that we cannot reject the null hypothesis that the synthetic and real images are drawn from the same distribution. The results confirm the quality of the models for mitochondria and nucleoli and the lower quality of the U-Net Golgi model.

Another question we want to address is if we can study the latent relationship between the shape and position of sub-cellular organelles. We fit a simple linear regression with organelles’ position as explanatory variables and the first principle component of their corresponding SPHARM descriptors as response, the result-in *R*^2^ scores are 0.0158, 0.0092, 0.08, and 0.0077 for nucleoli, Golgi apparatus, mitochondria, and desmosome, respectively., suggesting a lack of correlation between organelle’s shape and position.

## 4 Discussion

In this paper, we introduce new approaches for evaluating the quality of generative models of subcellular organelles. The first is based upon the expectation that those organelles are largely spatially-distinct from each other in the subcellular environment. To measure the extent to which generative models meet this expectation, we have introduced measures of exclusivity in which we compare the predictions for different proteins from the same transmitted light image and measure the overlap among different organelles at the same pixel position. High values of exclusivity were observed for a number of organelles. However, some organelles, such as microtubules, actin filaments and endoplasmic reticulum, showed fairly low values. Such low values can result from blurry predictions, but can also reflect similarity between the appearance of different organelles in light microscope images (which results in the models predicting that both organelles appear at the same position). Such ambiguity is certainly to be expected, and the ability of some organelles to be predicted with high exclusivity is a remarkable feature of the approach developed by Johnson *et al*. [2017]. Our criteria reflects which organelles are suited to that approach.

Given our finding that some organelle predictions overlapped significantly, we developed a retraining process that uses the predictions for other subcellular organelles to guide model learning for a particular organelle. This was done by adding an L1-loss term that penalizes overlaps, and the results demonstrated that organelle exclusivity could be improved without major loss in reconstruction error. Future work may focus on exploring other choices of loss function to model the interdependencies among channels. We anticipate that our approach to object-wise representations of cell images can not only be used to evaluate different generative modeling approaches, but also to enable better modeling of phenotypic changes in subcellular organization resulting from perturbations or genetic differences.

Our second evaluation approach is based on the fact that many organelles exist as discrete objects. We therefore asked whether the shape and spatial distributions of the objects in synthetic images adequately reproduce those of the objects in real images. Using a proven approach for shape modeling we demonstrated that generative models for at least some organelles do in fact produce objects similar to real images. To our knowledge this is the first demonstration that this is the case.

We have also introduced an improved GAN-based pipeline for learning to predict realistic fluorescence microscope images for subcellular organelles from bright-field microscopy images. The synthetic images produced by our new method are better predictions by all of our criteria. Our proposed approach on object-wise metrics has demonstrated the practicality and efficiency on evaluating the morphology of subcellular structures. A possible direction for future work may be to bring the spirit of object-wise metrics into the construction of learning objectives or as part of the feedback in the learning process.

## 1 Supplementary Tables and Figures

**Table S1.**
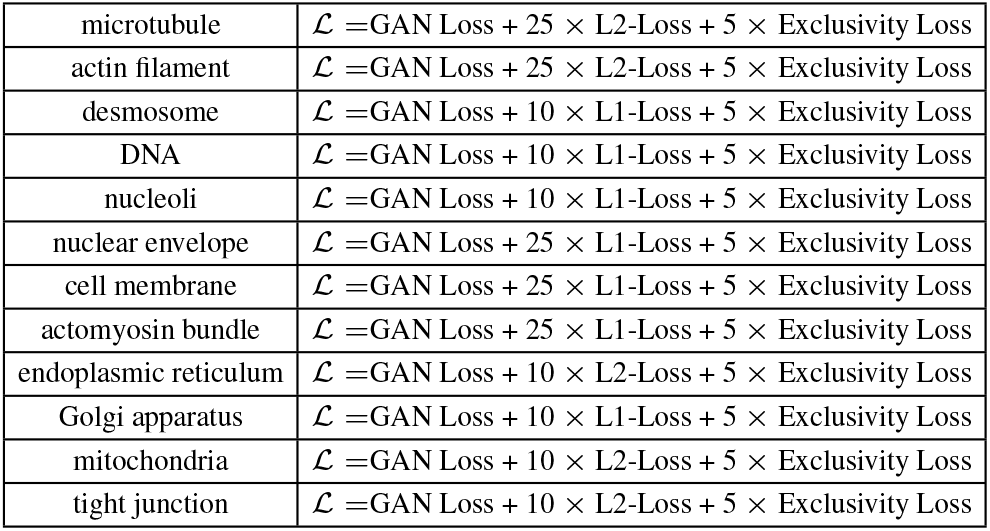
The loss functions used in retraining the Vox2Vox-RU model

|  |  |
| --- | --- |
| microtubule | $\mathcal{L} = \text{GAN Loss} + 25 \times \text{L2-Loss} + 5 \times \text{Exclusivity Loss}$ |
| actin filament | $\mathcal{L} = \text{GAN Loss} + 25 \times \text{L2-Loss} + 5 \times \text{Exclusivity Loss}$ |
| desmosome | $\mathcal{L} = \text{GAN Loss} + 10 \times \text{L1-Loss} + 5 \times \text{Exclusivity Loss}$ |
| DNA | $\mathcal{L} = \text{GAN Loss} + 10 \times \text{L1-Loss} + 5 \times \text{Exclusivity Loss}$ |
| nucleoli | $\mathcal{L} = \text{GAN Loss} + 10 \times \text{L1-Loss} + 5 \times \text{Exclusivity Loss}$ |
| nuclear envelope | $\mathcal{L} = \text{GAN Loss} + 25 \times \text{L1-Loss} + 5 \times \text{Exclusivity Loss}$ |
| cell membrane | $\mathcal{L} = \text{GAN Loss} + 25 \times \text{L1-Loss} + 5 \times \text{Exclusivity Loss}$ |
| actomyosin bundle | $\mathcal{L} = \text{GAN Loss} + 25 \times \text{L1-Loss} + 5 \times \text{Exclusivity Loss}$ |
| endoplasmic reticulum | $\mathcal{L} = \text{GAN Loss} + 10 \times \text{L2-Loss} + 5 \times \text{Exclusivity Loss}$ |
| Golgi apparatus | $\mathcal{L} = \text{GAN Loss} + 10 \times \text{L1-Loss} + 5 \times \text{Exclusivity Loss}$ |
| mitochondria | $\mathcal{L} = \text{GAN Loss} + 10 \times \text{L2-Loss} + 5 \times \text{Exclusivity Loss}$ |
| tight junction | $\mathcal{L} = \text{GAN Loss} + 10 \times \text{L2-Loss} + 5 \times \text{Exclusivity Loss}$ |

**Table S2.** The MSE on the test set of the initial and retrained U-Net and Vox2Vox RU models

|  | initial training |  |  | after retraining |  |
| --- | --- | --- | --- | --- | --- |
|  | U-Net | 3D Pix2Pix | Vox2Vox-RU | U-Net | Vox2Vox-RU |
| microtubule | 0.374 | 0.53 | 0.461 | 1.03 | 0.456 |
| actin filament | 0.439 | 0.596 | 0.606 | 0.763 | 0.662 |
| desmosome | 31.4 | 1.905 | 1.484 | - | 1.64 |
| DNA | 0.599 | 0.859 | 0.823 | 2.13 | 0.876 |
| nucleoli | 0.249 | 0.311 | 0.304 | 0.380 | 0.296 |
| nuclear envelope | 0.285 | 0.41 | 0.382 | 0.591 | 0.349 |
| cell membrane | 0.629 | 0.84 | 0.811 | 1.32 | 0.777 |
| actomyosin bundle | 0.767 | 1.027 | 0.942 | 0.855 | 1.37 |
| endoplasmic reticulum | 0.455 | 0.669 | 0.683 | 0.913 | 0.598 |
| Golgi apparatus | 0.882 | 1.476 | 1.40 | 0.948 | 1.76 |
| mitochondria | 0.513 | 0.722 | 0.620 | 0.550 | 0.697 |
| tight junction | 0.740 | 0.983 | 0.734 | 0.770 | 0.767 |
| Overall (no desmosome) | 0.5363 | 0.766 | 0.705 | 0.919 | 0.78 |
| Overall | 3.185 | 0.861 | 0.7691 | 3.535 | 0.856 |

**Table S3.** Number of subcellular objects found in the synthetic and real images. The average Hausdorff distance measures the quality of spherical harmonic transform modeling of the objects, lower is better.

| Model | U-Net | U-Net retrained | Vox2Vox-RU | Vox2Vox-RU retrained | Ground-truth |
| --- | --- | --- | --- | --- | --- |
| Mitochondria |  |  |  |  |  |
| # objects | 2538 | 2519 | 2659 | 2626 | 2591 |
| Average Hausdorff distance | 0.9985 | 1.0459 | 1.0152 | 1.087 | 1.0166 |
| Nucleoli |  |  |  |  |  |
| # objects | 357 | 349 | 416 | 380 | 386 |
| Average Hausdorff distance | 1.0439 | 1.0185 | 0.9951 | 1.0277 | 1.0136 |
| Golgi apparatus |  |  |  |  |  |
| # objects | 1156 | 1041 | 1508 | 1437 | 1211 |
| Average Hausdorff distance | 0.9122 | 0.9086 | 0.9484 | 0.9819 | 0.9522 |
| Desmosome |  |  |  |  |  |
| # objects | - | - | 629 | 867 | 751 |
| Average Hausdorff distance | - | - | 0.8670 | 0.8712 | 0.8658 |

**Table S4.** Divergence of objects in spherical harmonic descriptor shape space from real and synthetic images and their p-values.

| Model | U-Net | U-Net retrained | Vox2Vox-RU | Vox2Vox-RU retrained |
| --- | --- | --- | --- | --- |
| Mitochondria |  |  |  |  |
| Shape divergence | 0.0679 | 0.0698 | 0.0664 | 0.0678 |
| p-value | 0.4 | (0.24 | 0.56 | 0.52 |
| Nucleoli |  |  |  |  |
| Shape divergence | 0.0716 | 0.06 | 0.0545 | 0.0547 |
| p-value | 0.28 | 0.8 | 0.94 | 0.96 |
| Golgi apparatus |  |  |  |  |
| Shape divergence | 0.0713 | 0.0815 | 0.0684 | 0.0653 |
| p-value | 0.12 | 0.02 | 0.38 | 0.7 |
| Desmosome |  |  |  |  |
| Shape divergence | - | - | 0.0754 | 0.0748 |
| p-value | - | - | 0.16 | 0.14 |

**Table S5.** KL divergence in subcellular spatial distributions between real and synthetic images.

| Model | U-Net | U-Net retrained | Vox2Vox-RU | Vox2Vox-RU retrained |
| --- | --- | --- | --- | --- |
| Mitochondria |  |  |  |  |
| Spatial divergence | 0.0004 | 0.0008 | 0.0025 | 0.0011 |
| p-value | 0.98 | 0.8 | 0.06 | 0.38 |
| Nucleoli |  |  |  |  |
| Spatial divergence | 0.0007 | 0.0013 | 0.0005 | 0.0007 |
| p-value | 1 | 0.98 | 1 | 1 |
| Golgi apparatus |  |  |  |  |
| Spatial divergence | 0.0073 | 0.0053 | 0.0014 | 0.0029 |
| p-value | 0.02 | 0.04 | 0.98 | 0.4 |
| Desmosome |  |  |  |  |
| Spatial divergence | - | - | 0.1893 | 0.3045 |
| p-value | - | - | 0.02 | 0.02 |

**Fig. S1.**
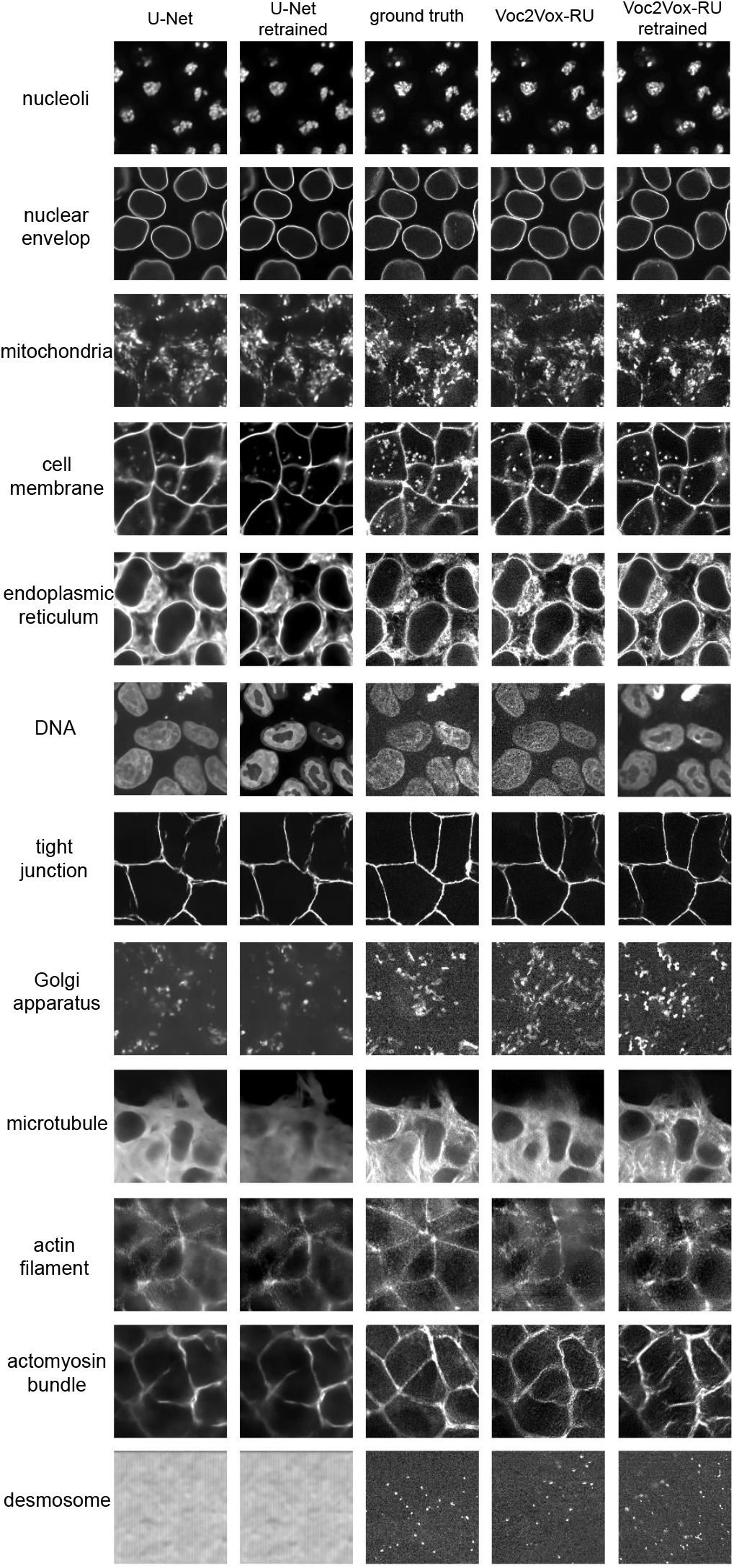
Example fluorescence tag predictions from U-Net (1st column), retrained U-Net (2nd column), real images (3rd column), and Vox2Vox-RU (4th column), and retrained Vox2Vox-RU (5th column). Note that desmosome is not retrained with U-Net, therefore the 1st column and 2nd column of the last row remain the same.

**Fig. S2.**
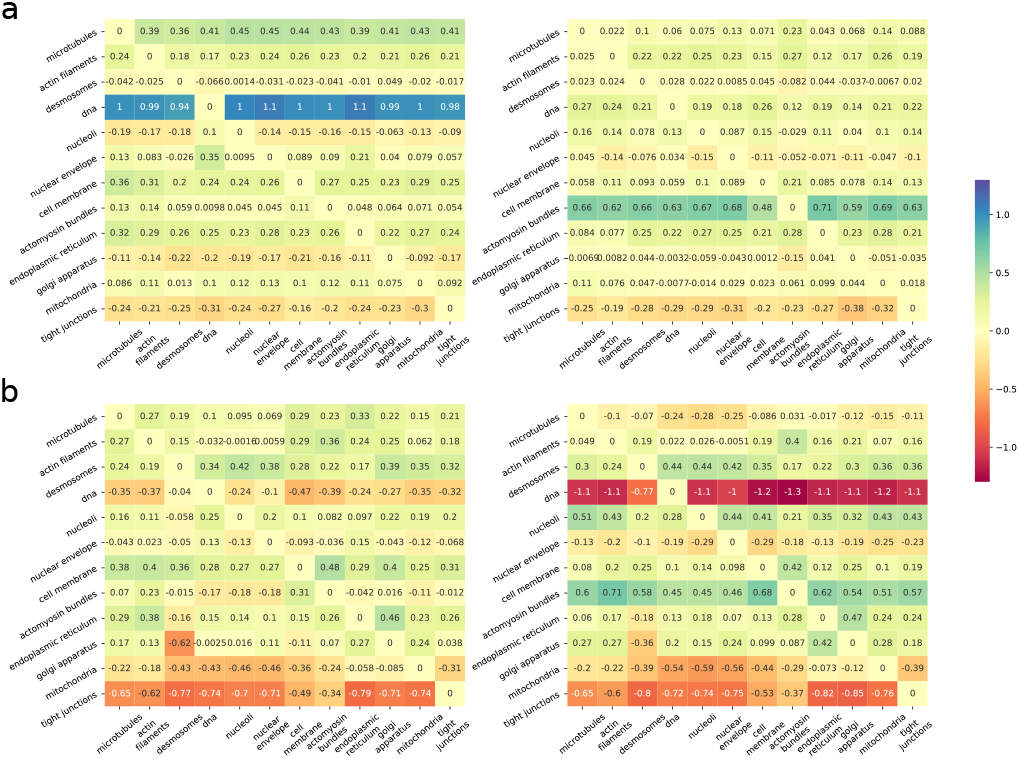
(a) The element-wise differences between the pairwise exclusivities of predictions from models before and after retraining of U-Net (left) and Vox2Vox-RU (right). (b) The element-wise differences between the pairwise exclusivities of predictions from Vox2Vox-RU and from U-Net models before (left) and after (right) retraining

**Fig. S3.**
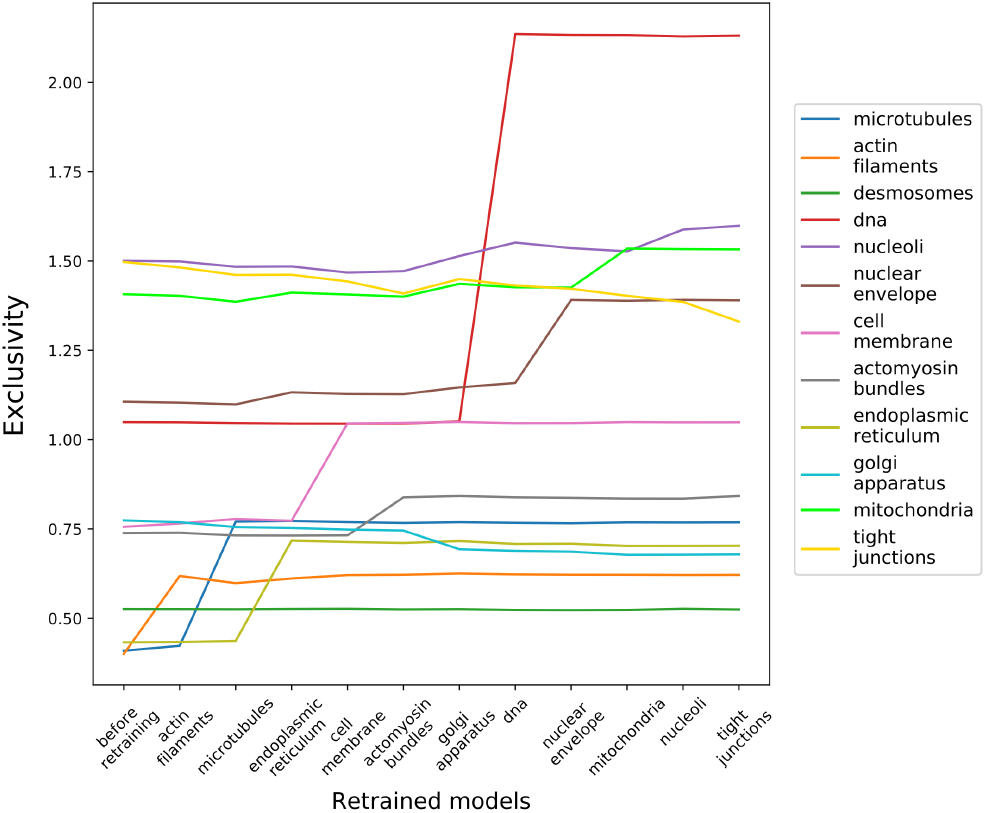
Individual organelle exclusivity changes during the retraining of U-Net models. The x-axis shows the organelle names in the retraining order and the lines with different colors show the individual organelle exclusivity changes.

**Fig. S4.**
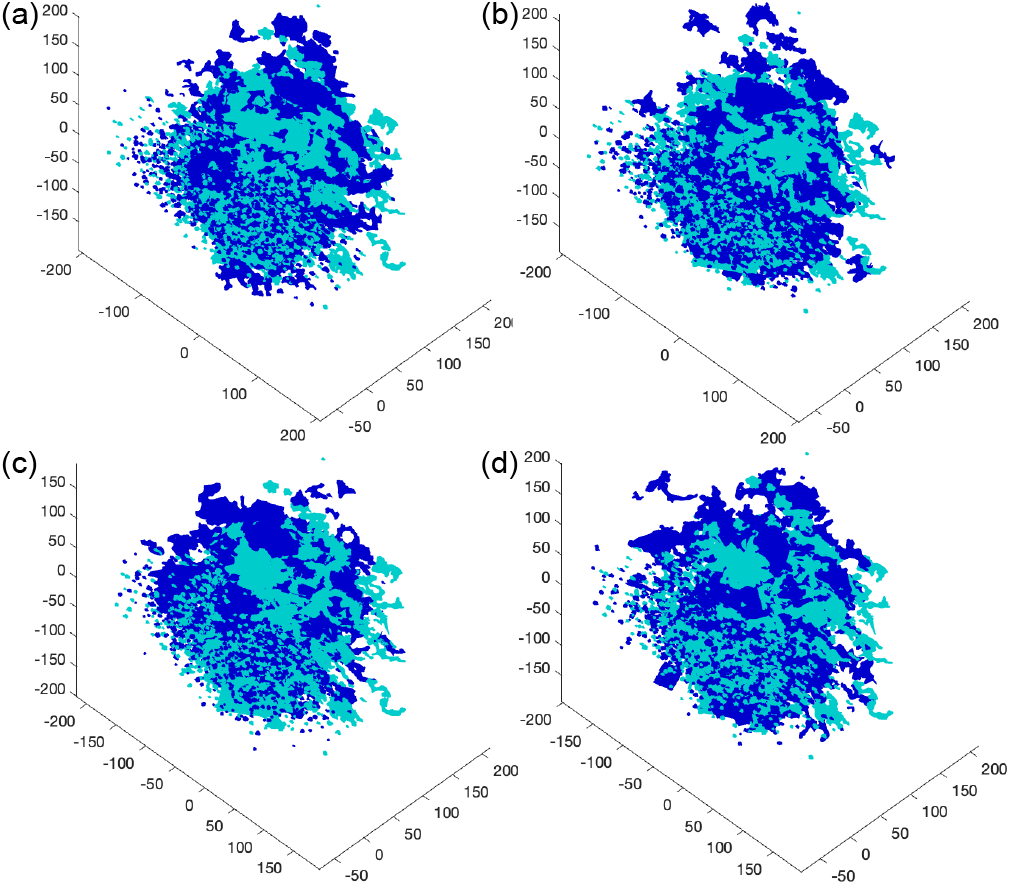
Visualization of mitochondria object shapes in the reduced spherical harmonic descriptor space. The light blue objects are from real images; the dark blue objects are from initial (a) and retrained (b) U-Net model without retraining and from initial (c) and retrained (d) Vox2Vox-RU model.

**Fig. S5.**
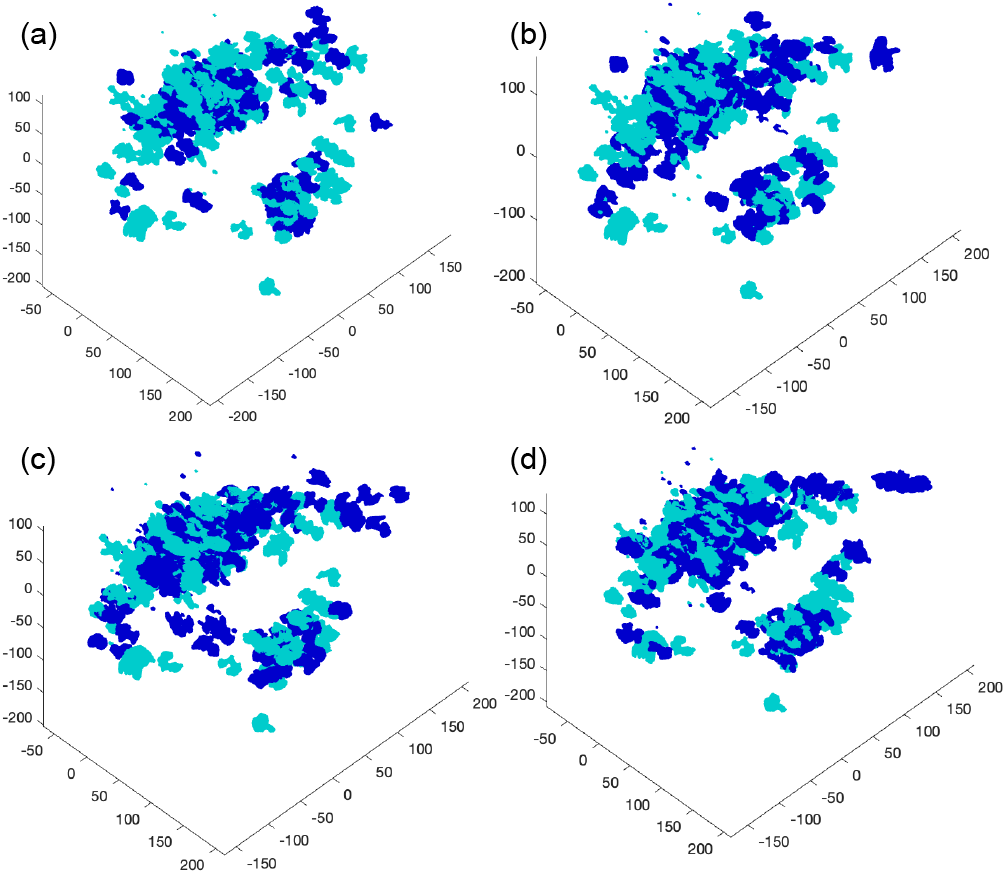
Visualization of nucleoli object shapes in the reduced spherical harmonic descriptor space. The light blue objects are from real images; the dark blue objects are from initial (a) and retrained (b) U-Net model without retraining and from initial (c) and retrained (d) Vox2Vox-RU model.

**Fig. S6.**
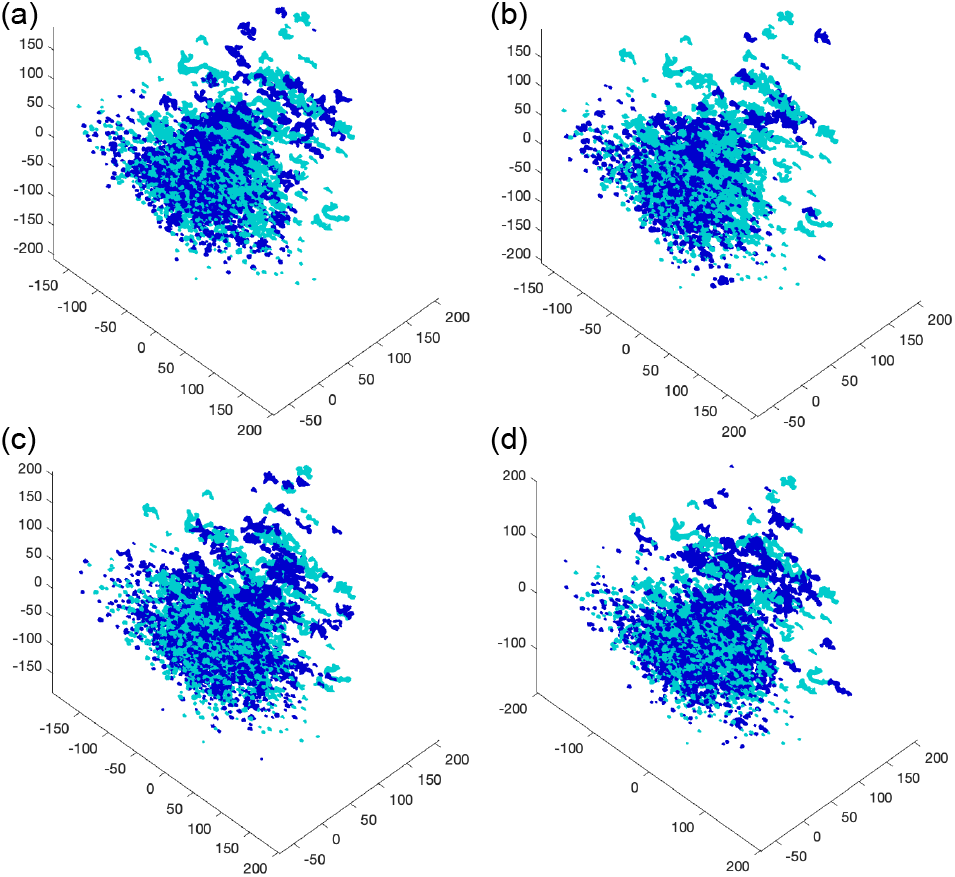
Visualization of Golgi apparatus object shapes in the reduced spherical harmonic descriptor space. The light blue objects are from real images; the dark blue objects are from initial (a) and retrained (b) U-Net model without retraining and from initial (c) and retrained (d) Vox2Vox-RU model.

**Fig. S7.**
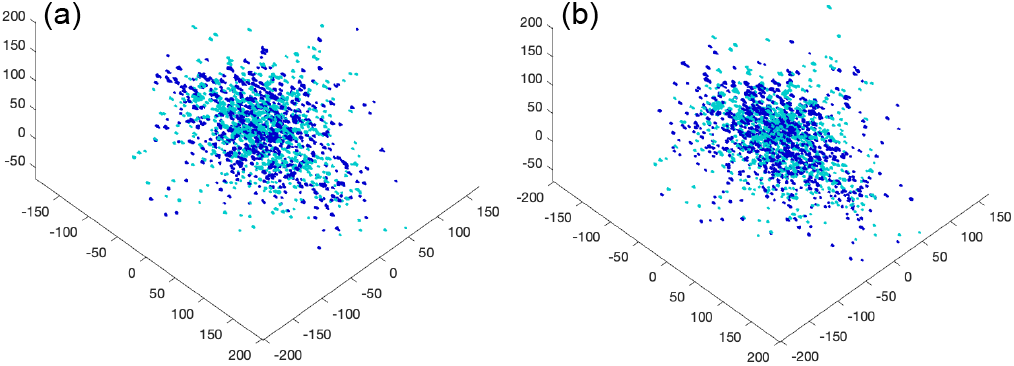
Visualization of desmosome object shapes in the reduced spherical harmonic descriptor space. The light blue objects are from real images; the dark blue objects are from initial (a) and retrained (b) Vox2Vox-RU model.Reduced spherical harmonic descriptor space of desmosome objects.

**Fig. S8.**
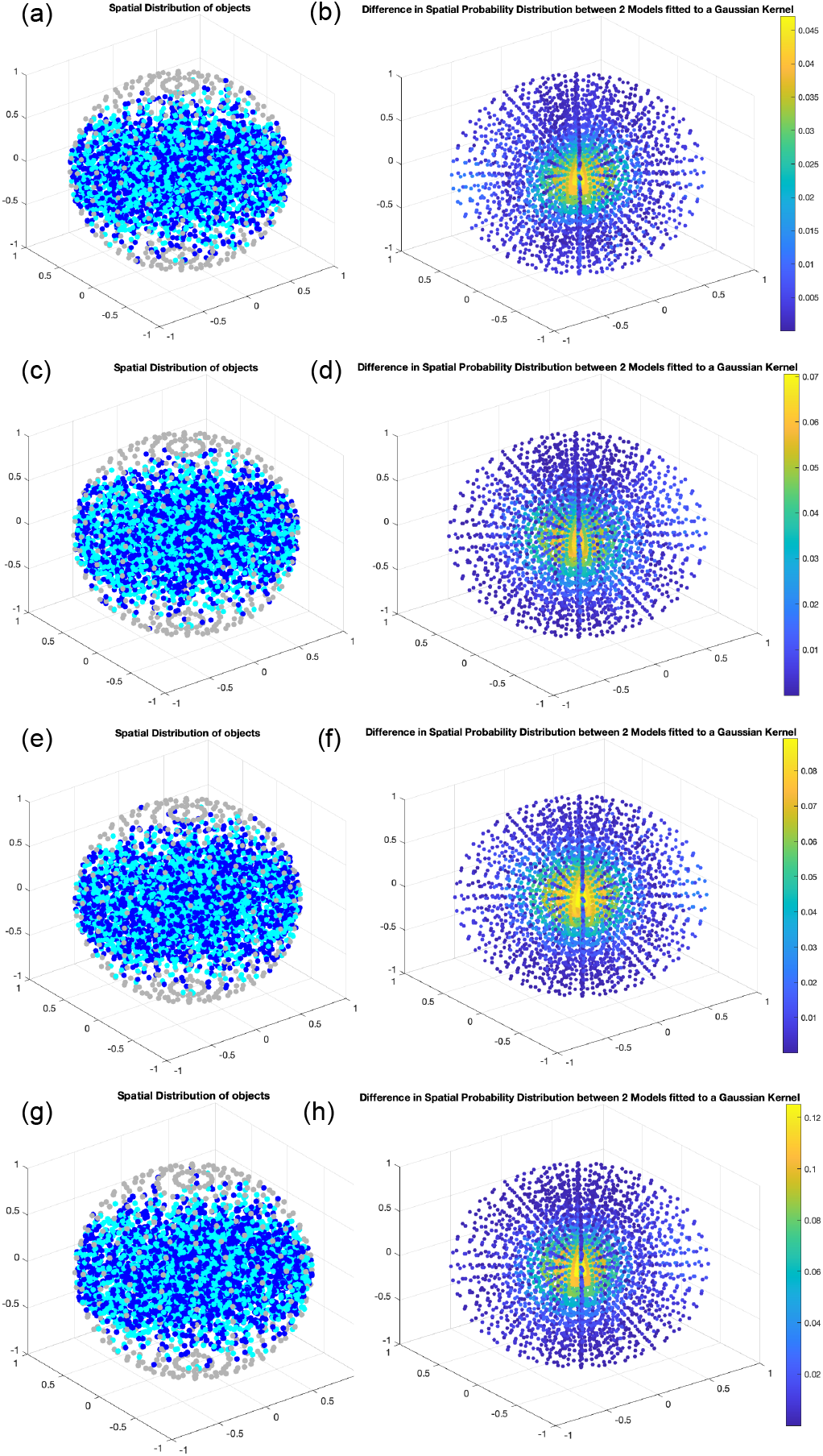
Spatial distribution analysis of mitochondria objects. Object positions are mapped onto a unit sphere whose center refers to the center of the cell and whose surface refers to the cell membrane. The light blue points are for objects from real images and the dark points refer to the objects from initial (a) and retrained (c) U-Net models and initial (e) and retrained (g) Vox2Vox-RU models. The absolute difference of the Gaussian smoothed spatial distributions between the objects from the real images and synthetic images are shown for the initial (b) and retrained (d) U-Net models and for the initial (f) and retrained (h) Vox2Vox-RU models.

**Fig. S9.**
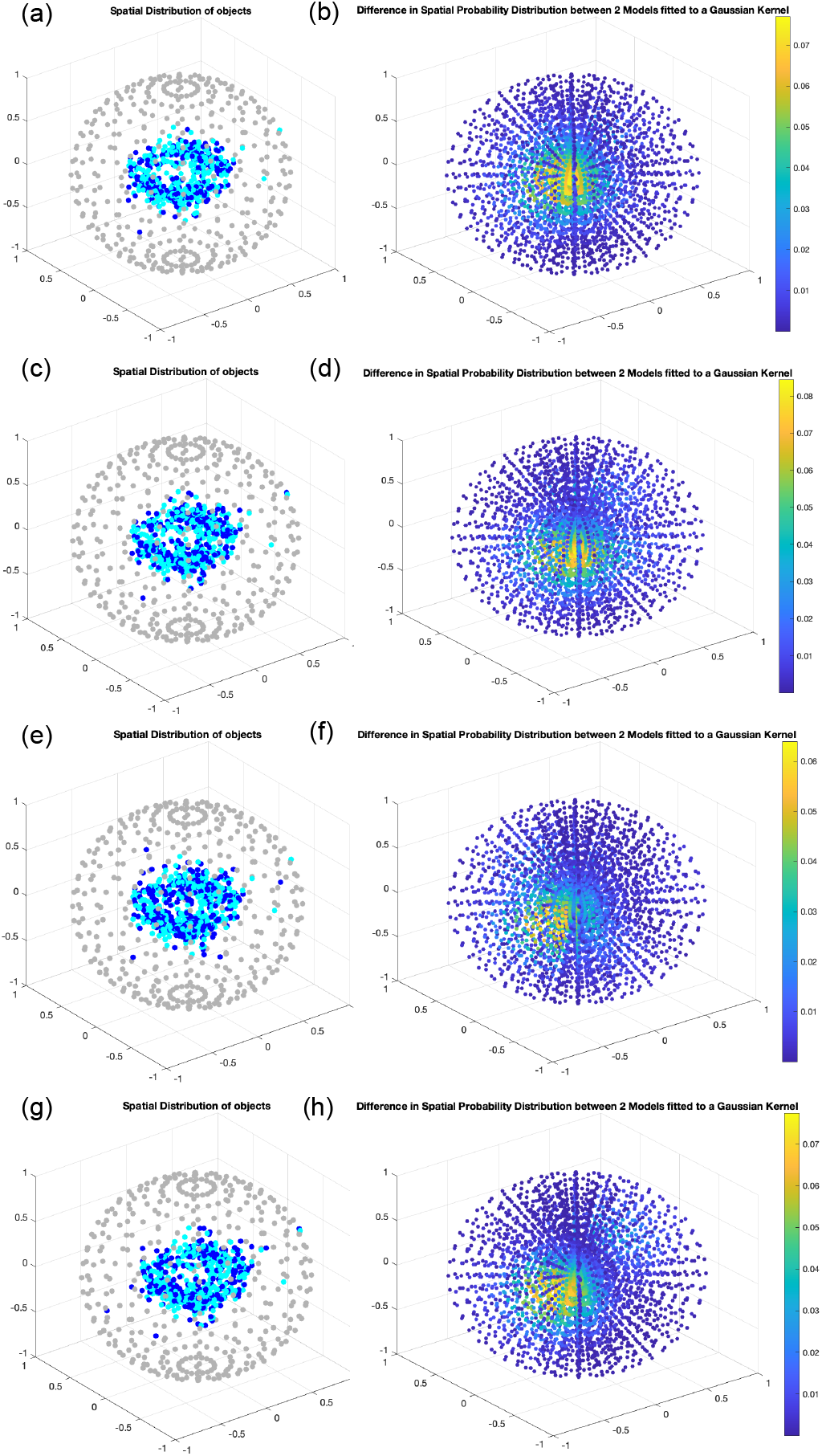
Spatial distribution analysis of nucleoli objects. Object positions are mapped onto a unit sphere whose center refers to the center of the cell and whose surface refers to the cell membrane. The light blue points are for objects from real images and the dark points refer to the objects from initial (a) and retrained (c) U-Net models and initial (e) and retrained (g) Vox2Vox-RU models. The absolute difference of the Gaussian smoothed spatial distributions between the objects from the real images and synthetic images are shown for the initial (b) and retrained (d) U-Net models and for the initial (f) and retrained (h) Vox2Vox-RU models.

**Fig. S10.**
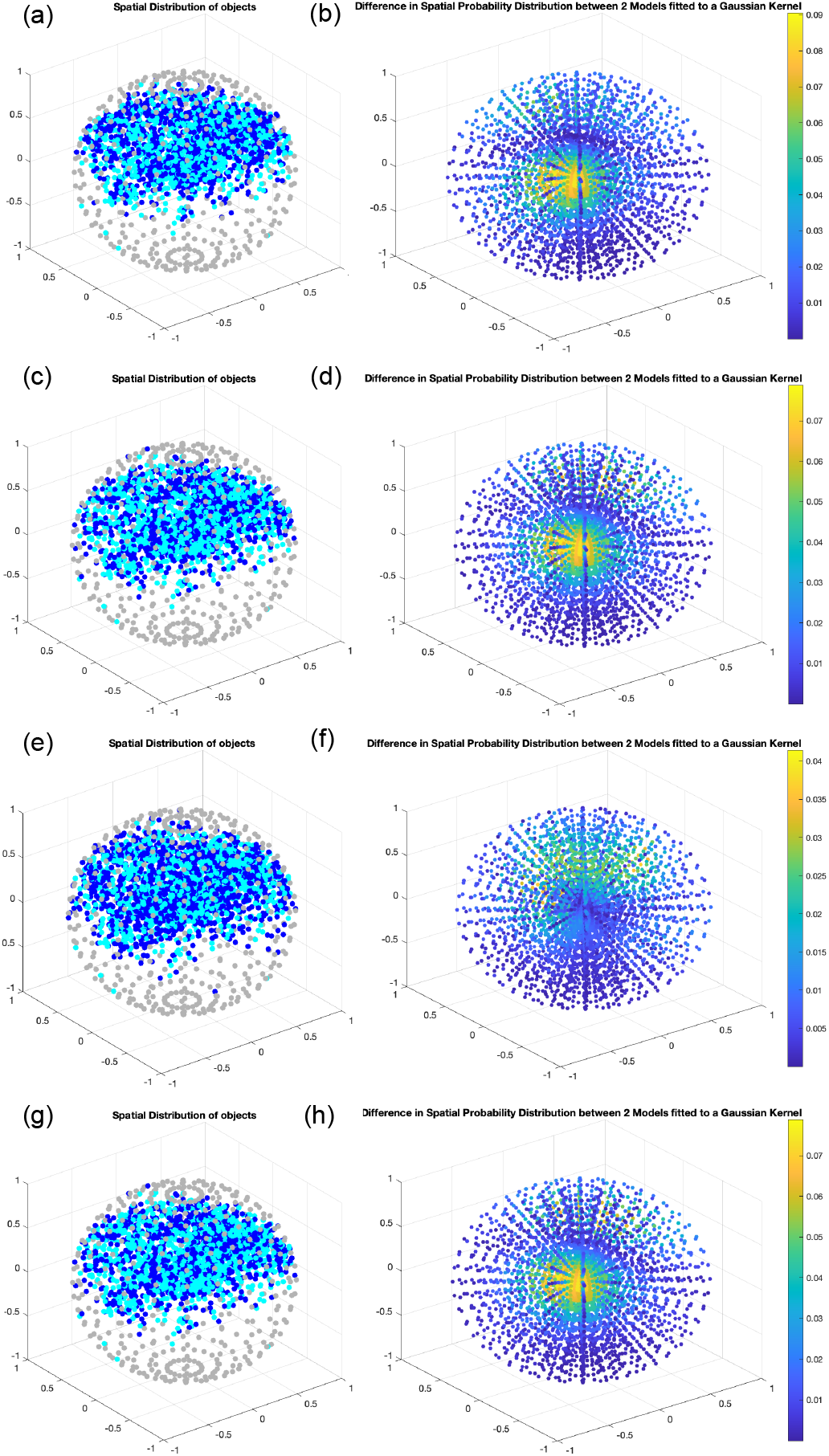
Spatial distribution analysis of Golgi apparatus objects. Object positions are mapped onto a unit sphere whose center refers to the center of the cell and whose surface refers to the cell membrane. The light blue points are for objects from real images and the dark points refer to the objects from initial (a) and retrained (c) U-Net models and initial (e) and retrained (g) Vox2Vox-RU models. The absolute difference of the Gaussian smoothed spatial distributions between the objects from the real images and synthetic images are shown for the initial (b) and retrained (d) U-Net models and for the initial (f) and retrained (h) Vox2Vox-RU models.

**Fig. S11.**
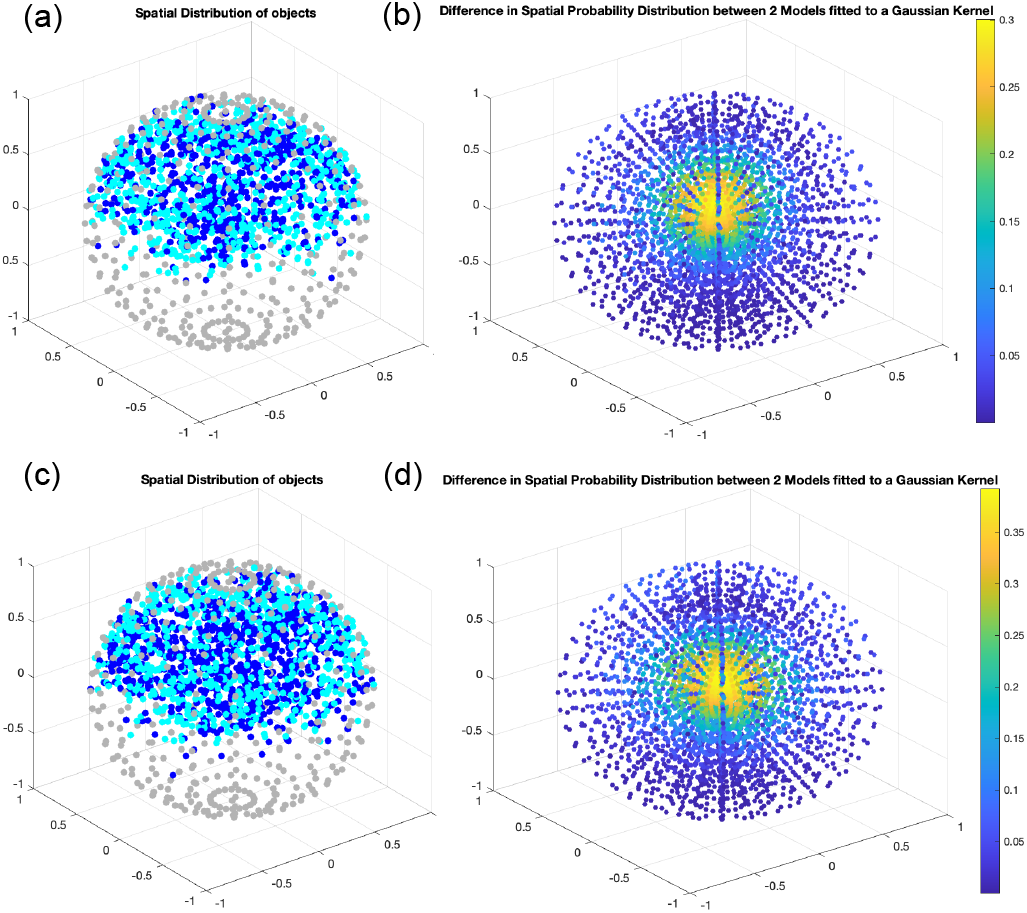
Spatial distribution analysis of desmosome objects. Object positions are mapped onto a unit sphere whose center refers to the center of the cell and whose surface refers to the cell membrane. The light blue points are for objects from real images and the dark points refer to the objects from initial (a) and retrained (c) Vox2Vox-RU models. The absolute difference of the Gaussian smoothed spatial distributions between the objects from the real images and synthetic images are shown for the initial (b) and retrained (d) Vox2Vox-RU models.

